# Agrin promotes coordinated therapeutic processes leading to improved cardiac repair in pigs

**DOI:** 10.1101/854372

**Authors:** Andrea Baehr, Kfir Baruch Umansky, Elad Bassat, Katharina Klett, Victoria Jurisch, Tarik Bozoglu, Nadja Hornaschewitz, Olga Solyanik, David Kain, Bartolo Ferrero, Renee Cohen-Rabi, Markus Krane, Clemens Cyran, Oliver Soehnlein, Karl Ludwig Laugwitz, Rabea Hinkel, Christian Kupatt, Eldad Tzahor

**Affiliations:** I. Medizinische Klinik & Poliklinik, University Clinic Rechts der Isar, Technical University Munich; DZHK (German Center for Cardiovascular Research), partner site Munich Heart Alliance Munich, Germany; The Department of Molecular Cell Biology, Weizmann Institute of Science, Rehovot, Israel; Department of Radiology, Klinikum Großhadern, LMU Munich; Institute for Cardiovascular Prevention (IPEK), LMU Munich; Department of Surgery, German Heart Center Munich; Deutsches Primatenzentrum GmbH, Leibniz-Institut für Primatenforschung, Department of Laboratory Animal Science, Göttingen, Germany

**Author notes:** both authors contributed equally (Andrea Baehr, Kfir Baruch Umansky). both authors share corresponding authorship Eldad Tzahor, Department of Molecular Cell Biology, Weizmann Institute of Science, Rehovot, Israel, Christian Kupatt, I. Medizinische Klinik & Poliklinik, Klinikum Rechts der Isar, Munich, Germany.

## Abstract

Ischemic heart diseases are classified among the leading cause of death and reduced life quality worldwide. Although revascularization strategies significantly reduce mortality after acute myocardial infarction (MI), a significant number of MI patients develop chronic heart failure over time. We have recently reported that a fragment of the extra cellular matrix (ECM) protein Agrin promotes cardiac regeneration following MI in adult mice. Here, we tested the therapeutic potential of Agrin in a preclinical porcine model, comprising either 3 or 28 days (d) reperfusion period. We first demonstrate that local (antegrade) delivery of recombinant human Agrin (rhAgrin) to the infarcted pig heart can target the affected regions in an efficient and clinically-relevant manner. Single dose of rhAgrin resulted in significant improvement in heart function, infarct size, fibrosis and adverse remodeling parameters 28 days post MI. Short-term MI experiment along with complementary murine MI studies revealed myocardial protection, improved angiogenesis, inflammatory suppression and cell cycle re-entry, as Agrin’s mechanisms of action. We conclude that a single dose of Agrin is capable of reducing ischemia reperfusion injury and improving cardiac function, demonstrating that Agrin could serve as a therapy for patients with acute MI and potentially heart failure.

## Introduction

In recent years, ischemic heart diseases have become the leading cause of mortality worldwide (1, 2). One of the most prevalent manifestations of ischemic heart disease is acute myocardial infarction (MI). In this injury, a coronary artery is occluded, in turn causing inflammation (3, 4), apoptotic, necroptotic and necrotic cell death (5–7), and formation of an akinetic fibrotic scar, triggering hypertrophy and fibrosis of the remote myocardium (adverse remodeling) (8, 9). These processes may lead to further deterioration of the heart culminating in chronic heart failure (CHF) (10). Therefore, preservation of viable heart muscle is of paramount priority in the current guidelines of the ACC/AHA (11) and the ESC (12), and is attempted via percutaneous coronary intervention or - in case of unavailability - thrombolysis. However, despite optimized treatment for acute MI, ischemic heart disease accounts for about half of the incidence of heart failure (13).

Unlike many other tissues, the adult heart in mammals and specifically adult cardiomyocytes are mostly post-mitotic (14), thus unable to divide to allow meaningful regeneration of the adult heart upon injuries. In contrast, the transient regenerative potential of the neonatal mouse heart is well established (15), and recent studies suggest a similar time window in neonatal pig hearts (16, 17), that could reflect humans (18). Successful reactivation of the neonatal regenerative program was shown by several groups as a mean to boost cardiac regeneration in adult mice (19–24).

In this context, we recently demonstrated that the ECM protein Agrin promotes heart regeneration in mice following MI (19). Agrin induces moderate cardiomyocyte cell cycle re-entry and division *in vitro* and *in vivo* and is required for the full regenerative capacity of neonatal hearts. *In vivo*, a single administration of recombinant rat Agrin (rrAgrin) promotes cardiac regeneration after injury. While the regenerative response of rrAgrin was partly attributed to cardiomyocyte proliferation, we suggested a pleiotropic mode of action, benefiting multiple repair processes.

Agrin is a large matrix proteoglycan known for its roles in neuromuscular junction (NMJ) development and function where it triggers the aggregation of acetylcholine receptors via the muscle-specific kinase (MuSK) (25, 26) and the low-density lipoprotein receptor related protein 4 (Lrp4) receptor complex (27). Agrin also binds and signals through α-Dystroglycan (Dag1) (28–30). In the heart, we demonstrated that Dag1 serves as Agrin receptor expressed by cardiomyocytes, mediating mild dedifferentiation and proliferation (19). The C’ terminus portion of Agrin, contains 3 laminin G-like (LG) domains, of which the LG1 and LG2 are essential for Dag1 binding, while LG3 is essential for MuSK-Lrp4 binding (31, 32).

Mouse MI models do not necessarily predict clinical outcomes in patients, due to differences in vessel structure, immune system as well as experimental features such as highly invasive thoracotomy, that are not clinically relevant. Therefore, examining the effect of a potential therapy in an MI model with patient-like instrumentation, dosage and application-venue may help to bridge the gap from bench to bedside. Accordingly, we designed a porcine MI study to test the distribution and efficacy of human recombinant Agrin. We found that intracoronary antegrade injection of Agrin is the most efficient delivery method for localized Agrin distribution in the injured heart. Using this approach, we observed that Agrin has a distinct cardio-protective effect in the subacute healing phase of infarcted hearts, and can potentially prevent heart remodeling and progression into heart failure. We could also show a conserved pleiotropic mechanism of action of recombinant Agrin in both mouse and pig MI models. Taken together, our findings support the application of recombinant Agrin as a novel therapy in patients with acute myocardial infarction, and potentially to prevent the onset of CHF.

## Results

### Local delivery method of rhAgrin into the ischemic region of the pig heart

In search for an efficient way to specifically deliver rhAgrin into the infarcted heart, we utilized an ischemia-reperfusion (I/R) model of MI in pigs (33, 34) (Figure 1A). The left anterior descending artery (LAD) was occluded for an hour, before reperfusion and injection of rhAgrin, delivered in one of three methods: antegrade infusion into the LAD, retroinfusion into the anterior interventricular vein (34) or intramyocardial delivery into the infarct border zone. After an additional 60 minutes, hearts were harvested, sectioned into 36 segments, and each segment was annotated as infarct (black area), border zone or remote myocardium (Figure 1B). To examine rhAgrin protein distribution, we extracted lysates from the different segments (Figure 1C). Levels of rhAgrin distribution in the heart was measured by western blot in 26 samples taken from 7 pig hearts. This analysis revealed that the antegrade delivery was the most efficient trajectory method (Figure 1D). Consistent with a regional distribution, we found the highest levels of rhAgrin in the infarct and border zone samples and almost none in the remote region of the heart (Figure 1D). Importantly, we could not detect rhAgrin in other tissue samples (lung, liver, spleen) collected from these animals (Supplemental Figure 1). Collectively, this experiment highlights the antegrade delivery as an efficient method that may be utilized for drug delivery to the infarct region of the pig heart.

**Figure 1.**
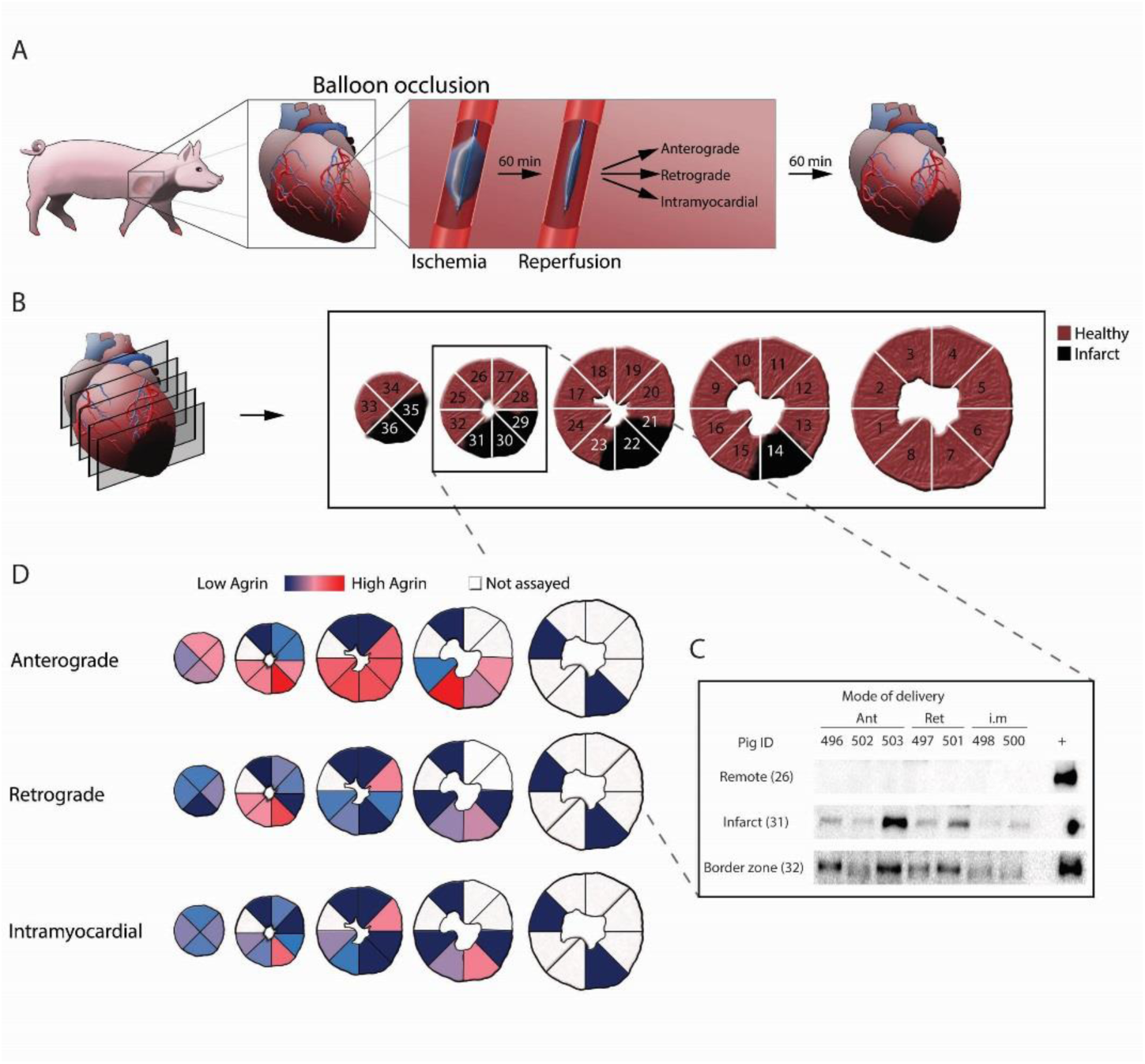
Comparison of rhAgrin delivery methods into the infarcted pig’s heart. (**A** and **B**) Schemes describing the delivery experiments: (**A**) Schematic description of the porcine MI ischemia-reperfusion model (60’ balloon occlusion). During reperfusion, rhAgrin (33µg/Kg) was administered in one of three trajectories: anterograde (n=3), retrograde (n=2) or intramyocardial (n=2) and hearts were harvested after an additional 60’. (**B**) Schematic description of the sectioning strategy used to sample infarct (in black), border zone and control (remote regions) of the infarcted hearts. (**C** and **D**) Assessment of Agrin distribution in the different segments of the infracted hearts. (**C**) Representative Western blot images comparing the amount of rhAgrin using all three trajectories in three segments: control (26), infarct (31), border zone (32). rhAgrin served as positive control (+); Ant-Antrograde, Ret-retrograde, i.m. - intramyocardial. (**D**) Color coded heat map of the average intensity at each segment for each trajectory (red-high Agrin, blue-no Agrin, White-not assayed). Each mode of delivery is presented in a different row.

### rhAgrin treatment promotes long-term improvement of cardiac function

We next set out to explore the effectiveness of rhAgrin treatment after MI in pigs. To do so, we treated infarcted pigs with rhAgrin (at 33μg/Kg) either by giving a single antegrade infusion immediately after reperfusion, or adding a second infusion 3 days post MI. Saline treatment was used as control (Figure 2A). Post-ischemic impairment of ejection fraction (EF, 24.78±1.02% (controls) was attenuated after rhAgrin administration (36.81±2.38%, single dose and 41.66±1.79%, dual dose, Figure 2, B and C). Further, rhAgrin treatment prevented left ventricular end-diastolic pressure (LVEDP) elevation, a prognostic hallmark of heart failure (17.72±0.82 mmHg in controls), again with no difference between one or two doses (10.26±0.36 mmHg, single dose and 10.13±0.92 mmHg, dual dose, Figure 2, D and E). Treatment of rhAgrin also resulted in increased stroke volume, as measured by MRI (Supplemental Figure 2). Because we could not detect significant differences between one or two doses of rhAgrin (Figure 2B-E), we concluded that a single dose of rhAgrin is sufficient to produce the beneficial effects. Importantly, both EF (Figure 2B) and LVEDP (Figure 2D) parameters revealed a short-term improvement already at 3 days post MI. Taken together, these findings suggest that rhAgrin treatment improves heart function post MI, with indications for a cardioprotective mechanism of action.

**Figure 2.**
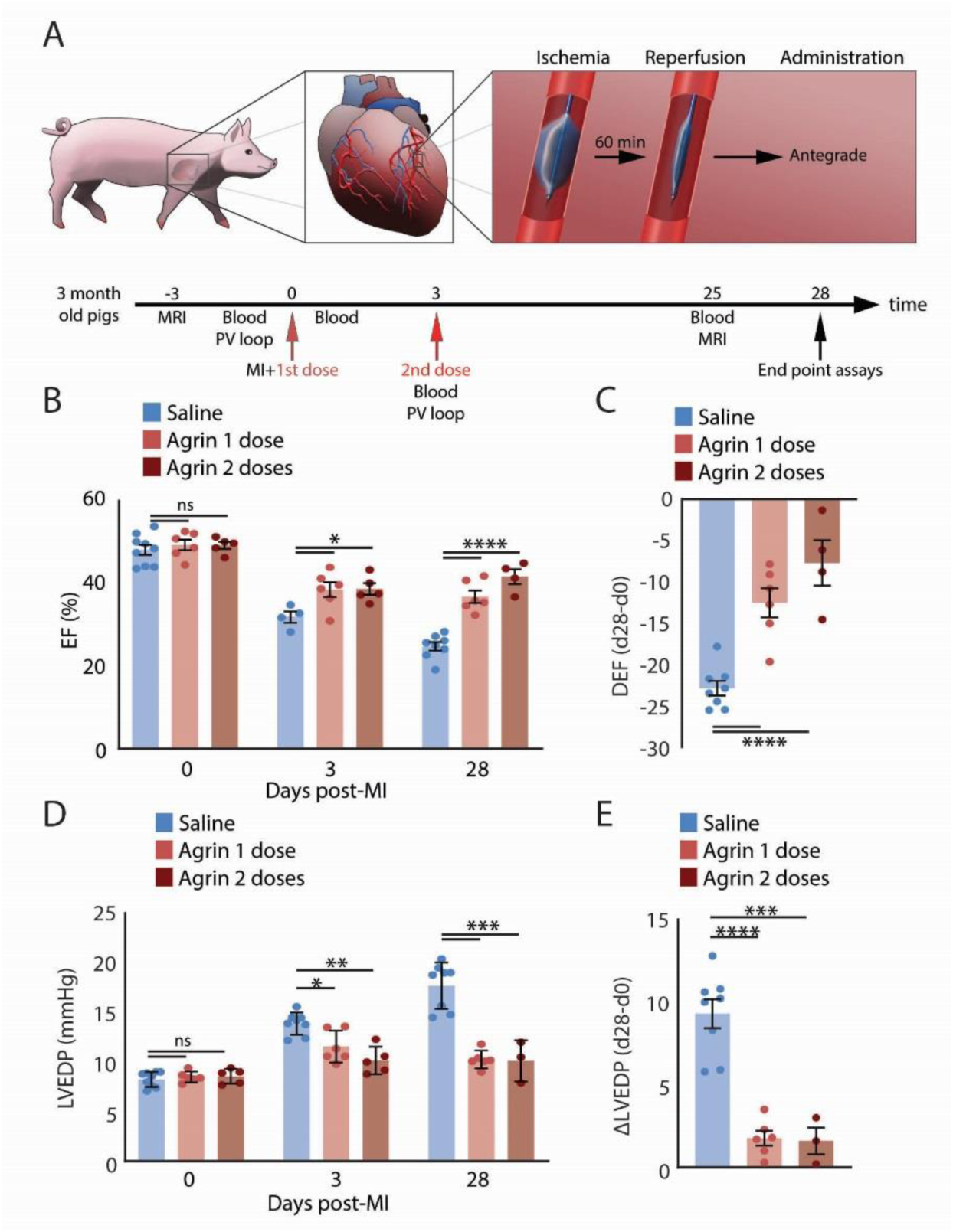
Heart function is improved in Agrin treated pigs post MI. (**A**) Scheme describing the experimental plan of the pig MI model. MI and Agrin treatment were applied as described in Figure 1A, using the antegrade method. Animals were treated with one or two doses of rhAgrin at day 0 and 3 of the experiment and followed for 28 days. Saline was used as control. Saline n=8, Single (1 dose) rhAgrin treatment n=6, Dual (2 doses) rhAgrin treatment n=5. (**B** and **C**) Bar graphs depicting the EF changes, derived from fluoroscopy analysis; (**B**) Bar graph depicting EF values at baseline, 3 and 28 days post MI. (**C**) Bar graph describing the reduction in EF in the different treatments at experimental end point. (**D** and **E)** Bar graphs showing the change LVEDP, as measured by PV-loop; (**D**) Bar graph depicting LVEDP values at baseline, 3 and 28 days post MI. (**E**) Bar graph demonstrating the changes in LVEDP in the different treatments at experimental end point; *-*p*<0.05, ***-*p*<0.001.

### rhAgrin reduces cardiac scarring and prevents remodeling of the infarcted heart

Prominent harmful consequences of MI are the excessive infarct scarring and adverse remodeling of the heart, which includes cardiomyocyte hypertrophy, ventricular dilation and increased heart weight. Therefore, we examined these parameters in the infarcted porcine hearts. In control pigs 28 days post MI, heart to body weight ratio (HW/BW) was increased from an average of 3.11±0.12 to 4.79±0.23g/kg, *p*<0.005, while rhAgrin treated hearts displayed a HW/BW ratio of 4.01±0.26g/Kg (single dose) and 3.77±0.32g/kg (dual dose) (*p*<0.05, compared to the control endpoint hearts Figure 3, A and B). Further, infarct size normalized to the left ventricular area was significantly reduced in rhAgrin treated hearts (from 21.03±2.16% to 10.39±1.65%), with a trend towards further reduction after 2 doses of rhAgrin (8.51±1.62%), as measured by TTC staining (Figure 3, C and D). This trend of reduced scarring was also detected using late enhancement MRI (Supplemental Figure 4, A and B). Of note, assessing the area at risk (AAR) at day 28 revealed similar area at risk in the different groups, with small variations within the groups (Supplemental Figure 3). This shows the strong reproducibility of the pig MI model and also implies that the hearts of both saline and rhAgrin treated animals suffered the same ischemic injury. Interstitial fibrosis was also reduced in the rhAgrin treated hearts, in the infarcted area and immediately adjacent to that (“infarcted area at risk”) (Figure 3, E and F) and border zones (Supplemental Figure 4, C and D). Cardiomyocyte hypertrophy was slightly reduced in rhAgrin treated hearts (Figure 3, G and H), which could suggest protective or even proliferative effects of rhAgrin on cardiomyocytes. Taken together, our data support the notion that rhAgrin prevents infarct scar expansion and remote myocardium remodeling post MI.

**Figure 3:**
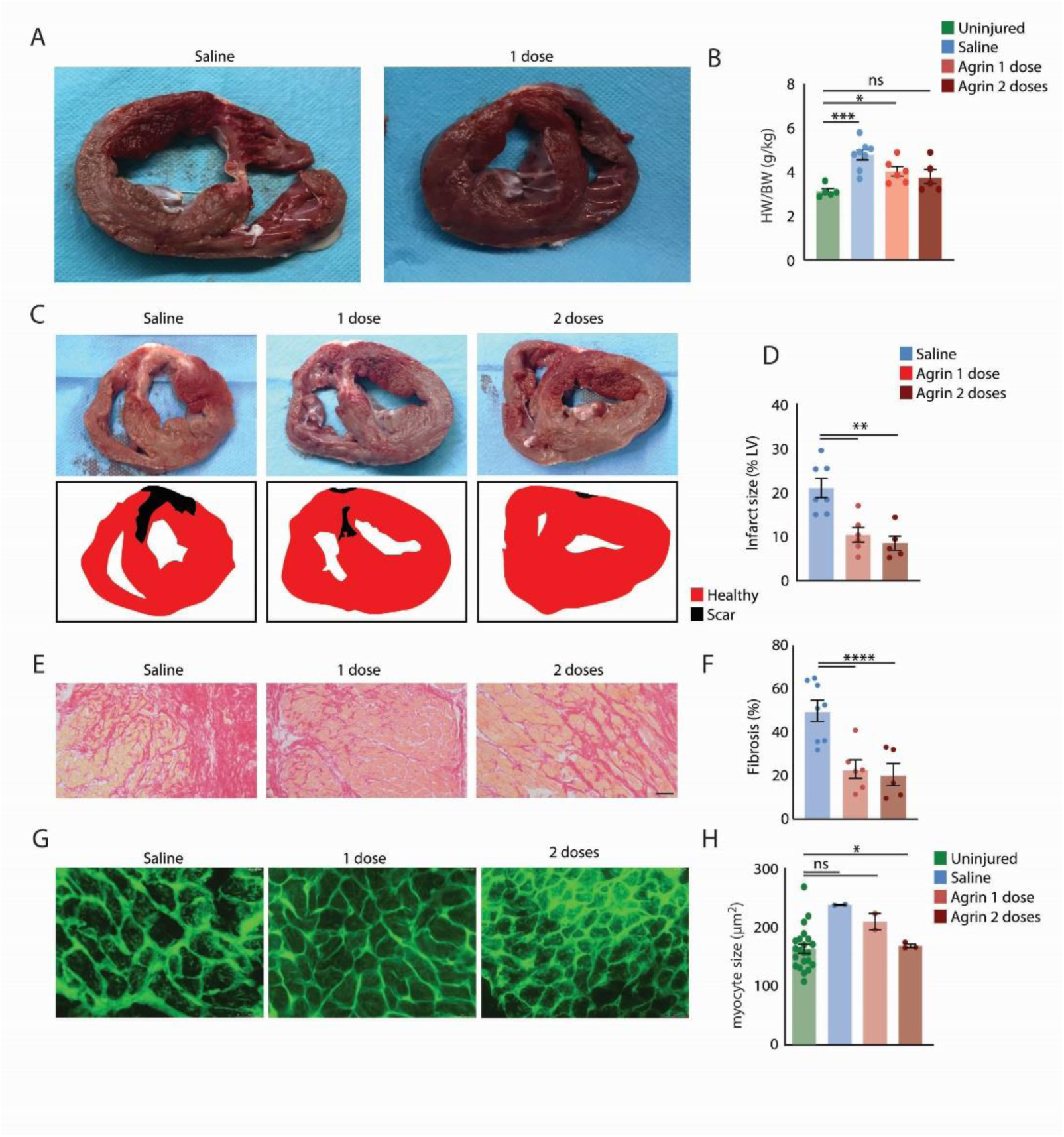
Heart remodeling is reduced in rhAgrin treated pigs post MI. (**A** and **B**) Heart weight to body weight ratio (HW/BW) (**A**) Representative images of saline and rhAgrin treated heart sections. (**B**) Bar graph showing the HW/BW at end stage. Age matched uninjured animals were used to establish baseline values; (**C** and **D)**. Scar assessment following Saline or rhAgrin treatment at 28 days after MI; (**C**) upper panel: representative images of heart sections after TTC staining (white represents scar tissue, red represents viable myocardium); lower panel: Same images with a graphic mask depicting healthy tissue (red) and scar (black). (**D**) Bar graph describing the scar tissue as a percent of the left ventricle wall, derived from TTC image analysis. (**E** and **F**) Interstitial fibrosis was measured in Saline/ rhAgrin treated hearts. (**E**) representative images of Sirius red stains of the infarcted area at risk. (**F**) Bar graph describing fibrosis as % of collagen in myocardial sections. (**G** and **H**) Assessment of CM size in the infarct zones of Saline/ rhAgrin treated infarcted hearts. (**G**) Representative images of heart sections stained with WGA; (**H**) Bar graph showing the differences in CM average size at the experimental end point. Age matched uninjured animals were used to establish baseline values; ns-non significant *-*p*<0.05, **-*p*<0.01, ****-*p*<0.0001.

**Figure 4.**
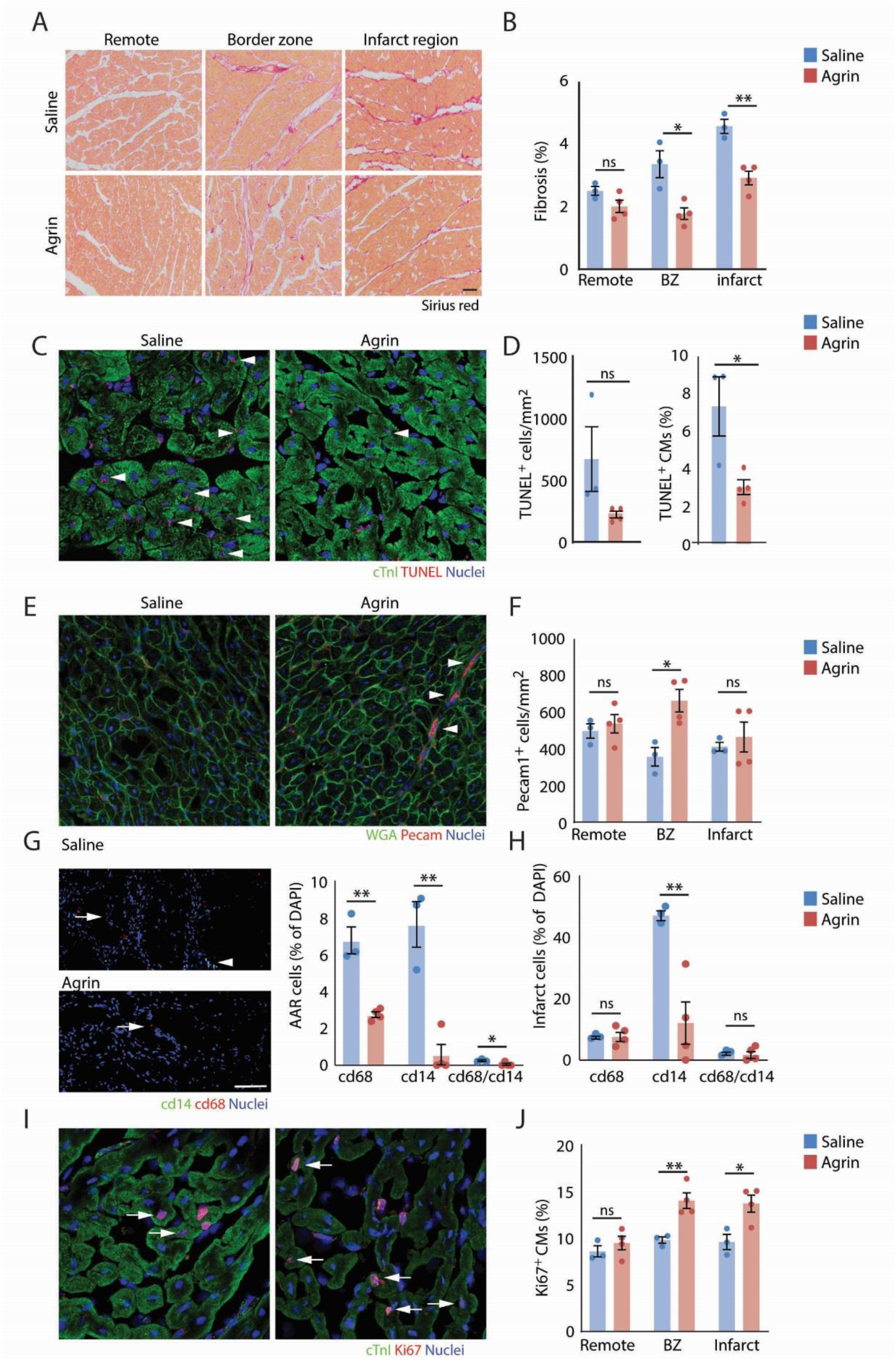
Agrin-induced repair mechanisms in the pig heart. Pigs underwent MI, and reperfusion was allowed for 3 days using the same monitoring strategy. At day 3, hearts were harvested and subjected to histological analysis. Saline n=3, rhAgrin n=4. (**A** and **B**) Interstitial fibrosis was assessed as in Figure 3E. (**A**) representative images of Sirius red from remote, border zone and infarct regions. (**B**) Bar graph describing fibrosis as % of collagen staining within myocardial sections. (**C** and **D**) Apoptotic cell death evaluated by TUNEL assay. cTnI IF was used as specific CM marker; (**C**) representative images of TUNEL and cTnI staining in border zone sections taken from Saline and rhAgrin treated hearts. White arrowheads highlight TUNEL positive cells; (**D**) Quantification of apoptotic cells (right panel). Bar graph depicting the number of apoptotic cells per myocardium area; (left panel) bar graph describing the number of apoptotic CMs. (**E**). Representative images of PECAM-1 staining in Saline and rhAgrin treated hearts border zone samples White arrowheads highlight PECAM-1 positive cells; (**F**) Bar graphs describing PECAM-1 positive cells in remote, border zone and infarct regions; **(G** and **H**) Macrophage IF analysis in the affected hearts by CD14 and CD68 staining (**G**). right panels: Representative images of CD14, CD68 and DAPI IF in border zone sections of Saline/ rhAgrin treated infarcted hearts; left panel: quantification of the CD14^+^, CD68^+^ and double positive cells in border zone sections (**H**) Bar graph describing the presence of CD14^+^, CD68^+^ and double positive cells in the infarct sections; (**I** and **J**) CM cell cycle re-entry/ proliferation was measured by Ki67 IF. cTnI IF was used as CM specific marker; (**I**) representative images of Ki67^+^ CM; (**J**) Bar graph describing Ki67 positive CM in the different regions of Saline/ Agrin treated hearts. *=*p*<0.05, **=*p*<0.001 vs. control.

### Conserved pleiotropic effects of Agrin treatment in both pig and mouse hearts

Given that our experiments indicated a prevention of adverse remodeling in post-MI hearts, we sought to analyze mechanisms which trigger infarct scar expansion, progressive loss of function and impairment of cardiac structure early after the ischemic insult. Hence, we assigned experimental groups to a 3d reperfusion period with or without rhAgrin treatment. Single dose administration of rhAgrin revealed a slight, but significant, improvement of systolic heart function (EF, Supplemental Figure 5A), with a tendency towards decreased LVEDP (Supplemental Figure 5B), similar to what we have previously measured in the chronic (long term) settings (Figure 2). No detectable difference in infarct size was evident following rhAgrin treatment (Supplemental Figure 5C). Nonetheless, rhAgrin treatment reduced interstitial fibrosis measured by Sirius Red staining (Figure 4, A and B), indicating a possible change in ECM composition. To examine a putative protective effect of rhAgrin, we stained for apoptotic cells using TUNEL. Indeed, rhAgrin reduced apoptotic cell death in both border zone and infarct regions (Figure 4, C and D), particularly in cardiomyocytes (Figure 4D). We also found higher number of PECAM-1^+^ capillaries in the border zone and the infarct zone in rhAgrin-treated hearts (Figure 4, E and F). This result suggests a protective role of rhAgrin in both endothelial and cardiomyocyte populations in the ischemic porcine heart. Moreover, the number of activated macrophages (CD68^+^ macrophage/monocyte marker (35)) and CD14^+^ inflammatory macrophage marker, (35)) was reduced in the rhAgrin-treated border zone sections (Figure 4G). A similar reduction in CD14^+^ cells was also noted in the infarct area of the rhAgrin treated hearts (Figure 4H). These results suggest that rhAgrin suppresses the inflammatory response that follows the MI. Finally, we could detect a significant increase in cardiomyocyte cell cycle re-entry measured by Ki67 staining in both border zone and infarct regions of the rhAgrin-treated pig hearts (Figure 4, I and J). Taken together, our results suggest that the regenerative effects of rhAgrin treatment in pigs include multiple target populations, and cellular processes.

**Figure 5.**
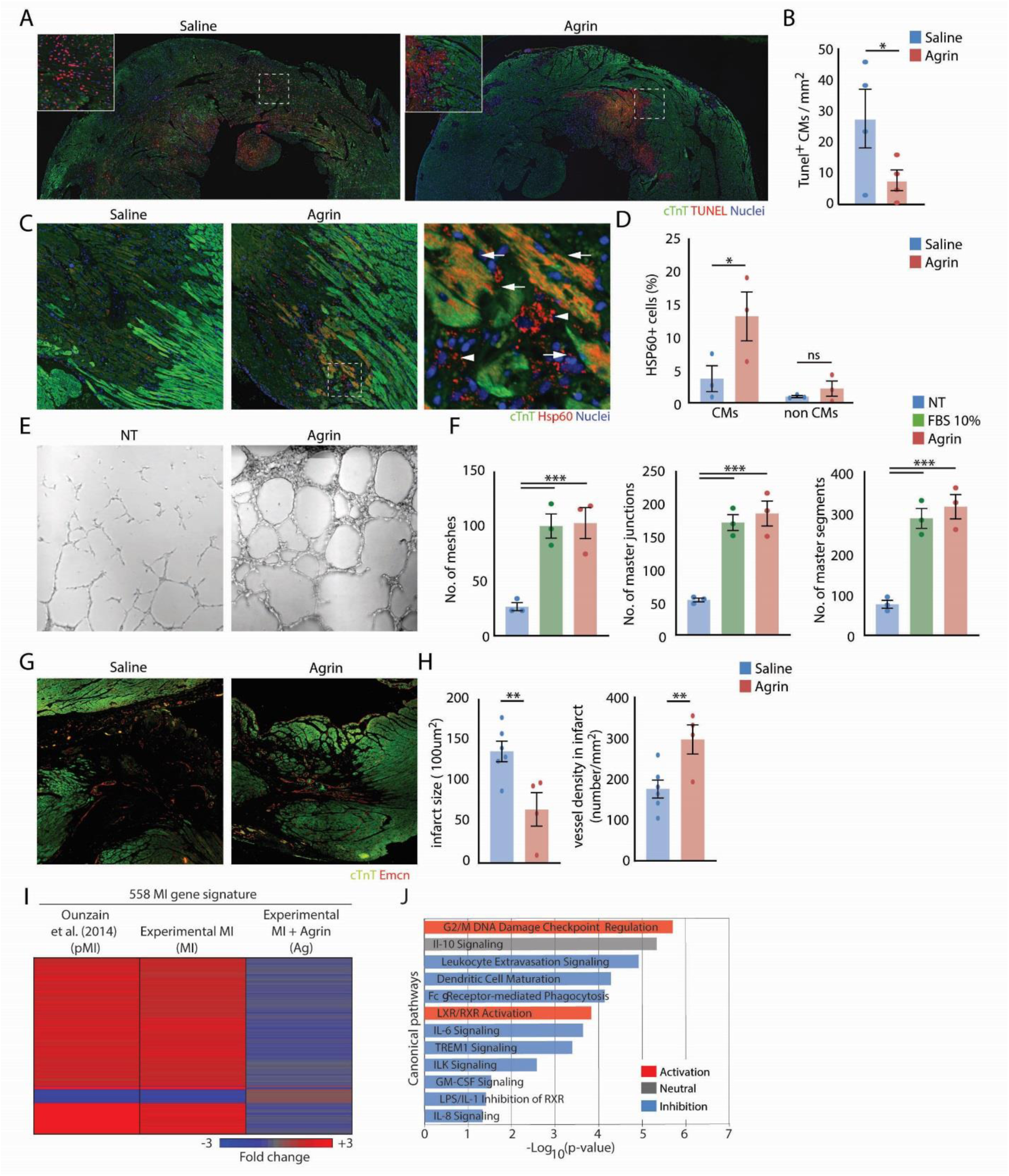
Conserved cellular mechanisms of Agrin treatment in the mouse MI model. Mice were subjected to MI, and treated with either Saline or rrAgrin. Hearts harvested for histological analysis at relevant time points. cTnT was used as CM marker. (**A** and **B**) Apoptosis was evaluated using TUNEL 24hr post MI, Saline n=4, rrAgrin n=4. (**A**) representative images of Saline/ rrAgrin treated hearts 24hr post MI; (**B**) bar graph describing apoptotic CM in Saline or rrAgrin treated hearts; (**C** and **D**) Hsp60 was imaged using IF in Saline/ rrAgrin treated mice, 2 days post MI. Saline n=3, Agrin n=3 (**C**) representative images of infarct and border zone regions, imaged for Hsp60; (**D**) bar graphs describing the Hsp60 positive cells in Saline/ rrAgrin hearts. (right panel) bar graph showing the Hsp60^+^ CM; (left panel) bar graph showing the Hsp60^+^ non-CM cells. (**E** and **F**) Assessment of rhAgrin induced *in vitro* tube formation in HUVEC cells. fetal bovine serum (FBS) was used as positive control, Saline was used as negative control. n=3 biological repeats for each group. (**E**) Representative images of HUVEC cultures 3 days post treatment with Saline (NT) or rhAgrin; (**F)** Quantification of different tube formation markers of Saline (NT), 10% FBS and rhAgrin; (**G** and **H**) Assessment of peri-infarct vasculature in MI mouse hearts, treated with PBS/ rrAgrin, 21 days post MI. Blood vessels were imaged by endomucin (Emcn) IF. Saline n=6, rrAgrin n=4 treated hearts. (**G**) representative images of Saline/ rrAgrin treated hearts; (**H**) bar graphs describing infarct size (right panel) and blood vessel density (left panel); (**I** and **J**) RNA – seq of infarcted and sham operated mouse hearts, treated with PBS/ rrAgrin. Hearts harvested 3 days post MI (Sham n=4, PBS n=4 and rrAgrin n=4). (**I**) Heat map depicting differentially expressed genes affected by Agrin. RNA-seq gene expression data was compared to an MI differentially expressed genes data base (38). Differentially expressed genes that showed similar pattern in our MI experimental setting (PBS vs. sham) and in (38) termed “MI signature”. The relative expression of these genes in (Ounzain, left panel) and our MI (Experimental MI, middle panel) and MI treated with Agrin (Agrin MI vs. PBS MI, Experimental MI+Agrin, right panel). (**J**) The genes shown in **I** were analyzed using ingenuity pathway analysis software. Prominent significantly enriched terms are shown for canonical pathways; (ns= non-significant, *=*p*<0.05, **=*p*<0.001, ***=*p*<0.001).

To better characterize these effects, we examined cellular events affected by recombinant rat Agrin (rrAgrin) in the adult mouse MI model (19). Ischemic hearts treated with rrAgrin showed reduced cardiomyocyte specific TUNEL staining 24hr post MI (Figure 5, A and B). In addition, the expression of Hsp60, a known anti-apoptotic marker in cardiomyocytes (36) was increased in rrAgrin treated hearts, 2 days post MI (Figure 5, C and D), underscoring a protective effect of rrAgrin in the mouse heart. Next, we examined the angiogenic role of Agrin in primary HUVEC cultures (37), and *in vivo* using infarcted mouse hearts (Figure 5E-I). In HUVEC, rhAgrin induced a strong tube formation response, similar to treatment with 10% fetal bovine serum (FBS) (Figure 5, E and F). Likewise, treatment of injured mouse hearts with rrAgrin showed an increase in blood vessel numbers 14 days post MI (Figure 5, G and H), specifically in small blood vessels (Supplemental Figure 6, A and B), indicating angiogenic effect of rrAgrin. Finally, we performed bulk RNA-seq on adult hearts following MI +/-rrAgrin (3 days post MI) and compared our findings to published MI RNA-seq dataset (38). This comparison allowed us to define common genes that are differentially expressed in infracted untreated hearts, serving as an “MI signature”. Of those, we identified 558 genes that were robustly repressed in the rrAgrin-treated hearts (Figure 5I). Bioinformatic analysis of the genes revealed a role for Agrin treatment in regulating multiple immune-related pathways (Figure 5J). Taken together, the data obtained from the mouse and pig models highlight Agrin’s pleiotropic effects, promoting cardiomyocyte protection, angiogenesis and attenuation of the inflammatory response, in addition to moderate cardiomyocyte proliferation. The conservation in the mechanisms of action in both mouse and pig injured hearts strengthen a pleotropic and broad cardiac repair program induced by Agrin treatment, making it highly relevant for treating human patients.

## Discussion

Perhaps one of the biggest challenges in the biomedical field, and particularly cardiology, is the gap from basic to clinical research arenas. More specifically, the translation of heart regenerative signals elucidated in rodents into large animal pre-clinical and clinical stages is missing. In this study we have made a step forward in this important path. We demonstrate an efficient and specific method of delivering rhAgrin into targeted regions of the infarcted heart in a preclinical pig model of acute MI, which can be easily and safely translated to patient treatment. Using the antegrade delivery method, we treated animals that underwent MI with rhAgrin. Similar to what we have previously found in the mouse model, rhAgrin improves both functional measurements and structural parameters in the pig heart. Functionally, besides actual EF improvement, rhAgrin almost completely prevented elevation of LVEDP post MI, a significant predictive indicator of heart failure. Structurally, the reduction in adverse remodeling, interstitial fibrosis and cardiomyocyte size, may suggest that rhAgrin treatment in acute MI setting could prevent the progression to heart failure.

Some of rhAgrin’s effects we observed in pigs could be detected as early as 3 days post MI, e.g., improved EF, reduced inflammation and apoptosis, indicating early cardioprotection - a beneficial outcome with regard to adverse remodeling. In line with this, the second dose of rhAgrin at 3 days post MI, had only marginal beneficial effects. Mechanistically, rhAgrin treatment in the first 3 days enhances myocardium salvage by preventing cell death, preservation of microvascular density, and by reducing the immune response. In addition to that, we observed initial evidence for cardiomyocyte cell cycle re-entry, as was already shown in the mouse model (19). Importantly, all cellular events elicited by rhAgrin in the pig heart were also evident in the mouse model, suggesting that Agrin effects are evolutionary conserved in mammals. Further, we could demonstrate that rhAgrin affected relevant human cultured cells, human iPS-derived cardiomyocytes (19) and HUVEC (this work) underscoring its translational potential in human patients.

The early (3d) positive effect of rhAgrin after acute MI is not caused by newly formed cardiomyocytes. Instead, we would argue that Agrin promotes cardiac repair in a pleiotropic manner, affecting several cell types – cardiomyocytes, endothelial, fibroblast and immune populations, and processes - cell survival, immune modulation and blood vessel survival/ renewal. Notably, a role for Agrin in promoting angiogenesis was demonstrated recently (39).

Previous studies addressing heart regeneration, dealing with either small or large animal models, utilize several delivery methods of therapeutic agents, many of which are clinically irrelevant. In several studies (including ours), animals were injected with compounds directly into the myocardium following thoracotomy (19, 40–42). Other studies used systemic administration methods, without looking for targeting the agent into the injured heart (43). Many of the recent clinical trials suffered from the inability to effectively deliver the desired agents into the myocardium, reviewed in (44). We set forth an empirically verified trajectory, namely antegrade infusion as an efficient and rather specific mode of delivery, which was previously found to be efficient for regional myocardial treatment with microRNA inhibitors (45). This is an established procedure in clinical trials, e.g. CUPID12 or Repair-MI (46), and can be easily adapted into the current PCI practice for treatment of acute MI.

We propose that Agrin exerts a pleiotropic therapeutic response, comprised of intrinsic (CM-specific) and extrinsic (immune cells, endothelial cells, fibroblasts (47)) arms that contribute to repair processes of injured animals. In line with this, it is becoming more evident that cardiac regeneration and repair strategies should focus on augmenting several cellular events (i.e. cardiomyocyte proliferation, angiogenesis, immune response modulation) (44).

We and others have previously demonstrated that activation of potent mitogenic signals could induce cardiac regeneration (20–22, 48–50). These mitogenic signals should be transient in nature, as demonstrated in a recent publication of prolonged treatment of infarcted pig hearts, leading to massive cardiomyocyte proliferation, which is deleterious and fatal due to arrhythmias (48). Therefore, it is important to design a transient regenerative signal, to ensure safety and prevent cardiotoxicity. A single dose, non-systemic rhAgrin treatment, combined with tissue clearance of about 2-3 days (in mice) (19), are in line with both transient and safety therapeutic considerations for treating heart diseases.

MI is the most common cause for heart failure (51) as many patients suffering from MI gradually deteriorate to heart failure. Male patients that survive the MI have a 3-fold increase in the risk of recurrent MI and 2-fold increase in risk of cardiovascular-related death within 3 years post-discharge. In women, these risks are even higher, 5.5- and 3-fold, respectively (52). Treatment with rhAgrin in pigs with severe MI improves EF and prevents LVEDP elevation and cardiac remodeling. These findings suggest that Agrin treatment for acute MI patients may serve as a preemptive therapy for heart failure.

This study sets forth Agrin as a putative therapy for MI in patients, including clinically-relevant protocol and conserved mechanistic insights. Our study demonstrates a consistent therapeutic role of Agrin in cardiac repair, with simple, safe and clinically relevant administration protocol, including initial dose and regiment protocol, which could be translated for patients. The full spectrum of Agrin molecular mechanisms, dose-range, toxicity and pharmacokinetics should be further investigated in a more rigorous manner. This study also highlights a growing need to introduce preclinical platforms in large animals to studies done in rodents. Taken together, our results underscore the potential of the matrix protein Agrin as a novel safe and simple therapy for patients with ischemic heart disease.

## Materials and Methods

### Animals

German pigs were purchased from a local farm. Animal care and all experimental procedures were performed in strict accordance with the German and National Institutes of Health animal legislation guidelines and were approved by the local animal care and use committees.

For the mouse experiments, we used adult (12 weeks) ICR female mice. All experiments were approved by the Animal Care and Use Committee of the Weizmann Institute of Science.

### Pig Ischemia/ reperfusion model

All pig experiments were conducted in the Institute for Surgical Research at the University of Munich. Pigs were anesthetized and instrumented as described previously (53). Briefly, a balloon was placed in the LAD distal to the bifurcation of the first diagonal branch and inflated with 6 atm. Correct localization of the coronary occlusion and patency of the first diagonal branch was ensured by injection of contrast agent. the percutaneous transluminal coronary angioplasty balloon was deflated after 60 minutes of ischemia; the onset of reperfusion was documented angiographically. At the onset of reperfusion, animals were treated with 33μg/Kg (in 5mL of saline) of rhAgrin (#6624-AG, R&D Biosystems) or saline alone as control.

For ejection fraction measurements, we obtained X-ray Fluoroscopy images, and analyzed them using ImageJ software. For LVEDP measurements, we used the PV-loop system (ADV500, Transonic, USA), with a pig tail catheter. Infarct size was assessed via methylene blue exclusion, tetrazolium red viability staining as described previously (54).

### Mouse MI model

Myocardial infarction was performed as previously described (19). Briefly, mice were sedated with isoflurane (Abbott Laboratories), followed by tracheal intubation and ventilation. Lateral thoracotomy at the third intercostal space was performed, and MI was induced by left anterior descending (LAD) coronary artery ligation, using 8-0 sutures. rrAgrin (50 μ l at 1 μ g per mouse, #550-AG, R&D biosystems) / PBS were injected intramyocardialy. Then, thoracic wall incisions were sutured with 6-0 non-absorbable silk sutures, and the wound was closed using skin adhesive. Mice were then warmed for several minutes until recovery.

### MRI measurement

#### Image acquisition

All subjects underwent cardiac magnetic resonance imaging (MRI) with a clinical 3-T MR scanner (Skyra; Siemens, Erlangen, Germany) by using a dedicated bogy phased-array coil. MRI was carried out before rhAgrin or placebo infusion, 3 days and 25 days weeks after MI. The MRI-protocol included scout, cine acquisitions in 2-, 4- and shot-axis views and contrast material-enhanced (0.2 mmol/kg Gd-DTPA (Magnevist, Bayer Schering Pharma, Berlin, Germany) LGE (late gadolinium enhancement) acquisitions in 2- and shot-axis views. During the study, ECG, heart rate, and respiration were monitored continuously. For the assessment of left ventricular volume, ejection fraction, myocardial mass, stroke volume and cardiac output ECG-gated, breath-hold, 2-D segmented fast low-angle shot (FLASH) cine images were obtained (field of view, 264 x 320 mm^2^; section thickness, 8 mm; image acquisition matrix, 240 x 158; band-width, 945 Hz/pixel; TR, 31.14 ms; TE, 1.52 ms; flip angle, 41°). To assess the size of myocardial infarct (MI) ECG-gated, breath-hold, single-shot gradient-echo pulse sequence were used (field of view, 285 x 380 mm^2^; section thickness, 8 mm; image acquisition matrix, 256 x 124; band-width, 465 Hz/pixel; TR 487.57 ms; TE 1.52 ms; flip angle 20°)(55).

#### Image Analysis

Image analysis were performed by one independent radiologist with 5 years’ experience. Quantitative evaluation of the MI size was performed by using a cardiac segmentation toolbox implemented as a plugin in the open-source OsiriX software(56). Following parameters were generated: MI area (cm^2^), MI weight (g), MI volume (mL) and MI percentage. Left ventricular volume (end systolic and end diastolic), ejection fraction, myocardial mass, stroke volume and cardiac output were calculated using an implemented plugin MRHeart in the OsiriX MD software (v.9.0., Pixmeo, Switzerland).

### Immunofluorescence

For TUNEL assay in the mouse and pig, we used In Situ Cell Death Detection Kit, TMR red (Sigma Aldrich, 12156792910) according to the manufacturer protocol. For mice IF, heart sections underwent deparaffinization and microwave antigen retrieval in EDTA or citric acid buffer, followed by gradual chilling. Samples were permeabilized with 0.5% Triton X-100 in PBS for 5 min and blocked with 5% bovine serum albumin (BSA) in PBS containing 0.1% Triton for 1 h at room temperature. Then, samples were incubated overnight at 4°C with the following antibodies diluted in 3% BSA blocking solution and 1% horse serum: Anti-cTnT (Abcam, ab33589), anti-cTni (Abcam, ab47003) were used to identify CMs. Anti-EMCN antibody [V.7C7.1] (ab106100, Abcam) was used to identify endothelial cells. Anti-Hsp60 (ab46798, Abcam) was used to identify the Hsp60 protein. After three 10-minute washes with PBS, samples were stained for 1 h at room temperature with relevant fluorescent labeled secondary antibodies (Abcam) followed by 10 min of DAPI staining for nucleus counter stain. Slides were mounted with Immu-mount (9990412, Thermo Scientific) and viewed under a fluorescence microscope (Nikon Intensilight Nikon eclipse 90i, Nikon or Olympus live cell imaging microscope) or spinning-disc confocal microscope (Carl Zeiss).

For pig IF, heart samples were frozen on dry ice, embedded in OCT and sectioned on to slides. Heart slides were fixed in cold Acetone, washed in PBS (5 min. 3 washes) and blocked with 10% NFS, for 60 min. Then, slides were incubated overnight at 4°C with the following antibodies (diluted in blocking buffer): Anti PECAM-1 (sc-376764, Santa Cruz, 1:50), anti Ki67 (AB9260, Millipore, 1:100), anti-cTnT (Abcam, ab33589, 1:200). The slides were washed with PBS (5 min,3 washes), stained with the relevant secondary antibody for 2 hr at RT. Slides wer mounted and visualized as above described.

### *In vitro* tube formation

For HUVEC tube formation, we used awell established protocol, described in (37). In short, we have thawed HUVEC cells and passage them to a new plate when they reach ~75% confluency. These cells were then plated, in 12 wells plates, at 1e5/ well, and grown overnight. Cells were then pre-conditioned in starvation (EBM-2, #00190860, Lonza) containing the relevant treatment for 16 hr. HUVEC were then trypsinied and moved to growth factor reduced matrigel covered 24 wells plates, and were grown in starvation+ treatment containing media for 8 hr. Cells were then PFA fixed, visualized and tube formation was measured by the HUVEC analysis tool for ImageJ.

### Bulk RNA-seq

For RNA-seq, adults (3 months) mice were subjected to LAD ligation or sham operation (as described in (19)). The LAD ligated animals were injected with either Agrin or PBS (vehicle) epimyocardialy immediately after MI. Hearts were collected 3 days post treatment, and RNA samples were purified using the miRNeasy kit (#1038703, Qiagen) according to the manufacturer instructions. Trueseq libraries were made and barcoded, and subsequently subjected to 60bp single read solexa illumine sequencing, using the Solexa HiSeq. For RNA analysis, Adapters were trimmed using the cutadapt tool (57). Following adapter removal, reads that were shorter than 40 nucleotides were discarded (cutadapt option –m 40). Reads were sub-sampled randomly to have 23M Reads. TopHat (v2.0.10) was used to align the reads to the mouse genome (mm10) (58). Counting reads on mm10 RefSeq genes (downloaded from igenomes) was done with HTSeq-count (version 0.6.1p1) (59). Differential expression analysis was performed using DESeq2 (1.6.3) (59, 60). Raw P values were adjusted for multiple testing using the procedure of Benjamini and Hochberg.

### Statistical Analysis

The results are given as mean±SEM. Statistical analysis were performed by 1-way ANOVA. Whenever a significant effect was obtained with ANOVA, multiple-comparison tests between the groups with the Student-Newman-Keuls procedure were performed (SPSS 20.0 statistical program or Graphpad).

## Author contributions

A.B., R.H., K.K., N.H., M.K. and K.B.U. performed the pig experiments under the guidance of C.K, M.K. and K.L.L. V.J., N.H., K.K., F.B., O.S. and T.B. performed the histological analysis of the pig experiments. O.S. performed the MRI analysis under the guidance of C.C. E.B., K.B.U, R.C.R and D.K. performed the mice *in vivo* experiments under the guidance of E.T. who also conceived this project. E.T, C.K, and K.B.U wrote the manuscript.

## Acknowledgments

We thank E.T. and C.K. laboratory members, O. Goresh and B. Siani for animal husbandry, and R. Levine, S. Goldsmith, C. Raanan and M. Osin for histology. **Funding**: This work was supported by grants to E.T. from the European Research Council (ERC StG #281289, CM turnover, and by the European Union’s Horizon 2020 research and innovation program, ERC AdG #788194, CardHeal), from the Britain-Israel Research and Academic Exchange (BIRAX), from Foundation LeDucq Transatlantic Network of Excellence, the Israel Science Foundation (ISF) and the Israel Ministry of Science & Technology. C.K. received funds from the BMBF (DZHK Large animal platform), the DFG (Transregional program SFB 267) and the Else-Kröner-Fresenius Foundation.

