## Supplemental figures for "Agrin promotes coordinated therapeutic processes leading to improved cardiac repair in pigs"

**Supplementary Materials:**

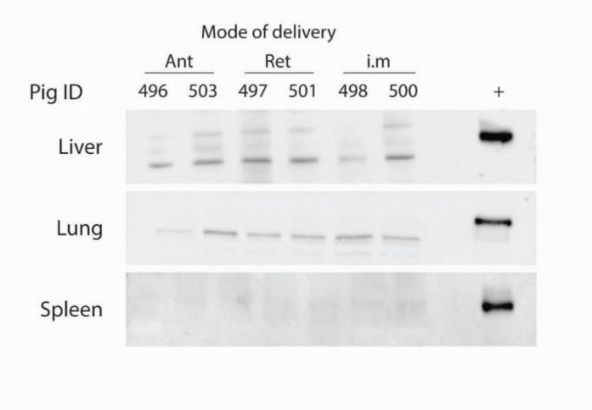


**Supplemental Figure 1. rhAgrin distribution in pig tissues.** Assessment of rhAgrin presence in Spleen, liver and lung samples, collected from the experiment described in Figure 1. Agrin amount was analyzed as described in Figure 1. Resulting western blot is shown.

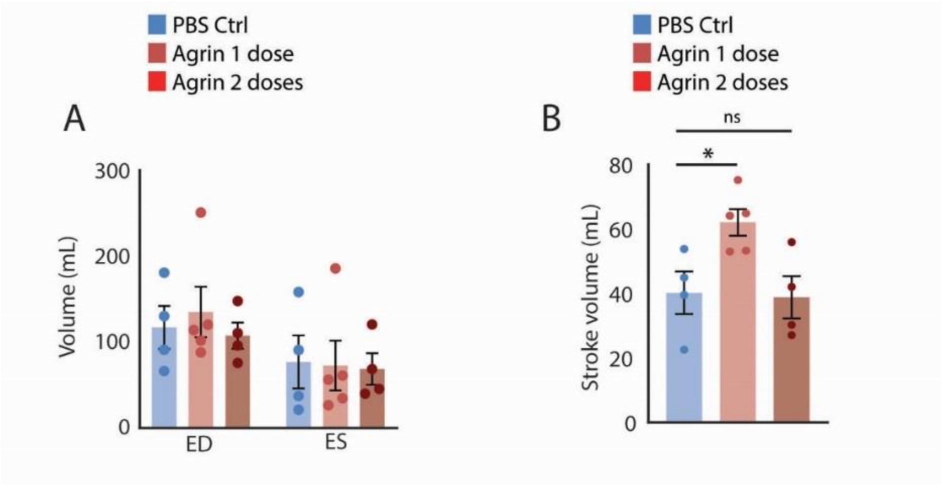


**Supplemental Figure 2. rhAgrin influences cardiac volumes.** Measurement of left ventricle volumes of Saline/ rhAgrin treated hearts 25 days post MI (MRI). (A) bar graph describing the changes in left ventricle volumes; (B) bar graph demonstrating the changes in stroke volumes at end point.

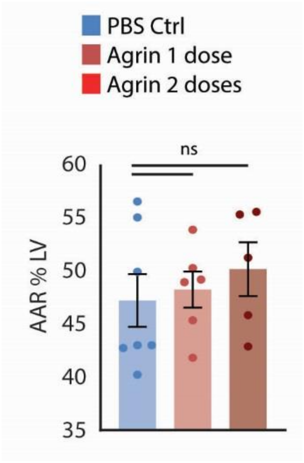


**Supplemental Figure 3. AAR size is not influenced by rhAgrin treatment.** Bar graph depicting the area at risk (AAR) as percent of left ventricle wall. The AAR was measured by applying TTC to the occlusion site in the LAD, and measuring the perfused area. AAR was similar in all groups, indicating similar LAD occlusions.

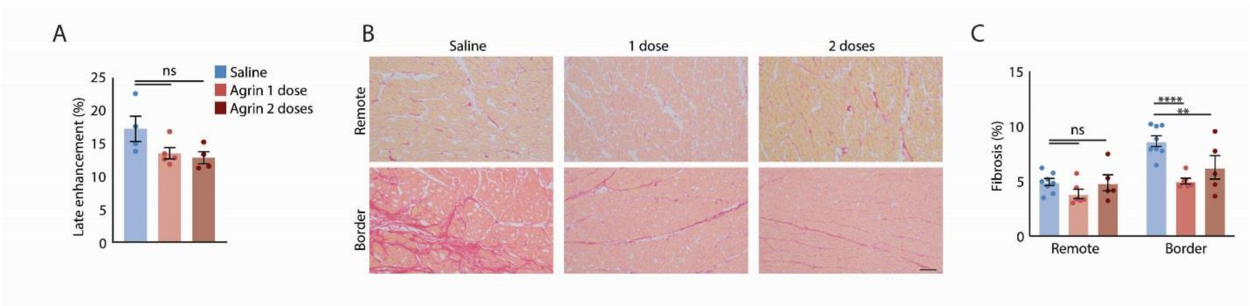


**Supplemental Figure 4. rhAgrin affects scar size and interstitial fibrosis.** (A) Assessment of Scar tissue volume by MRI. Scar was measured by late enhancement, 25 days post MI (as % of LV volume); (B and C) measurement of interstitial fibrosis in remote and border zones of injured hearts 28d post MI, as in Figure 3E. (B) representative Sirius Red images of remote and border zone in Saline, 1dose and 2 doses of Agrin; (C) quantification of interstitial fibrosis as % of collagen in myocardial sections; (ns= non-significant, **=$p$<0.001, ****=$p$<0.0001).

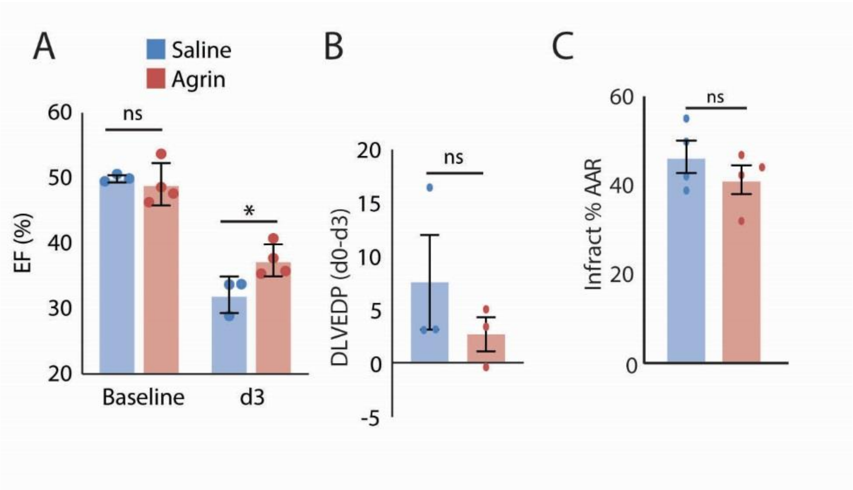


**Supplemental Figure 5. rhAgrin improves heart function improves 3 days post MI.** Assessment of rhAgrin effect on functional and structural cardiac parameters 3 days post MI: bar graphs demonstrating the mild improvement in EF (A) and LVEDP (B), and scar size (C) after 3 days perfusion experiments. N=3 for Saline, n=4 for rhAgrin.

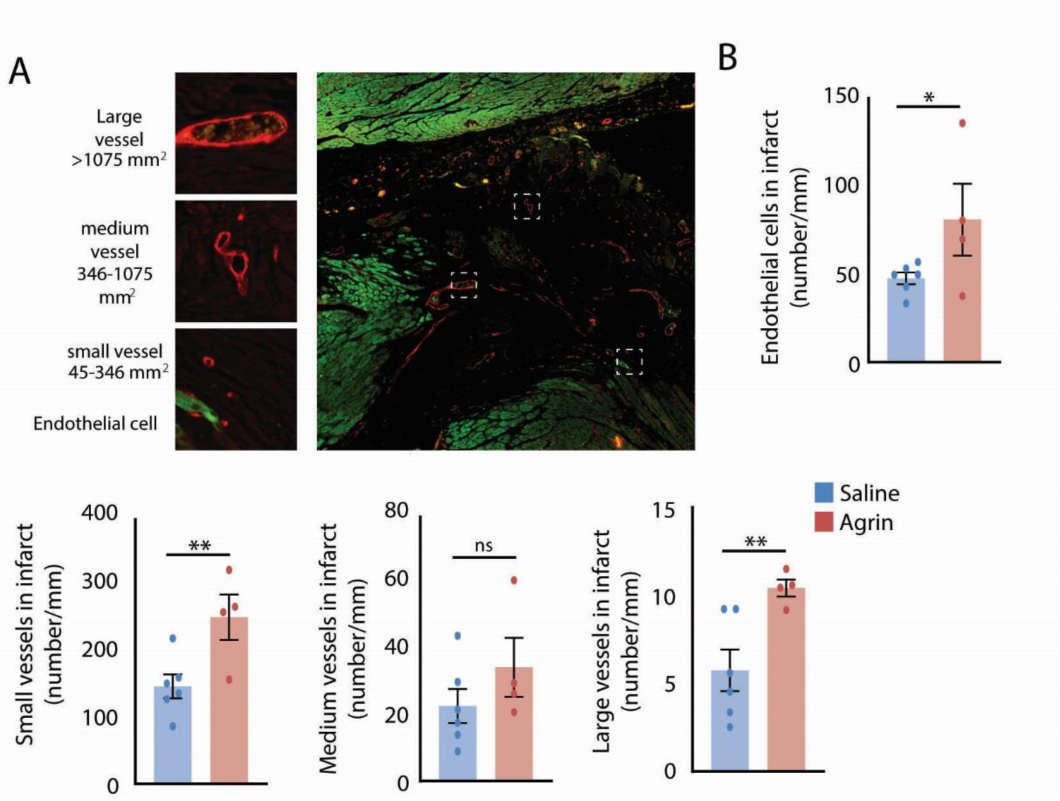


**Supplemental Figure 6. rrAgrin improves post MI micro vasculature in mice.** (A) reprehensive image of Ecmn IF in infarcted heart, 14 days post MI. The smaller right panels depict exmaples for large, medium and small blood vessels; (B) Bar graphs describing the differences in several vasculature parameters between the Saline and rrAgrin treated mice, including EC numbers in infarcted areas, and numbers of large, medium and small vessels.
